# Engineered Cas9 complexes establish an experimentally grounded benchmark for heterogeneous cryoEM reconstruction methods

**DOI:** 10.64898/2026.05.04.721978

**Authors:** Andrew V. Grassetti, Laurel F. Kinman, Joseph H. Davis

**Affiliations:** Department of Biology, Massachusetts Institute of Technology, Cambridge, MA; Department of Biochemistry and Biophysics, University of California, San Francisco, CA

## Abstract

Single-particle cryoEM is increasingly used to resolve conformational and compositional ensembles, yet objective evaluation of heterogeneous reconstruction methods remains limited by the scarcity of experimental benchmarks with per-particle ground-truth labels. Indeed, many widely used experimental”benchmark” datasets necessarily validate observed states retrospectively while purely synthetic datasets provide ground-truth labels but typically fail to capture experimentally realistic complexities including confounding structural heterogeneity, imaging noise, contaminants, and orientation biases, which dominate real-world analyses. Here we develop an experimentally grounded benchmark dataset for heterogeneous reconstruction using catalytically inactive *Streptococcus pyogenes* Cas9 bound to a constant sgRNA and to target DNA duplexes engineered to carry extensions of defined length. We assembled, purified, vitrified, and imaged thirteen complexes independently, such that the dataset-of-origin provides an unambiguous label for each particle’s encoded state while preserving the full experimental complexity of cryoEM data. Independent refinements of the pure datasets recover the engineered DNA-extension signal and define a simple quantitative readout, DNA-extension occupancy, that increases monotonically with designed extension length. The same reconstructions also reveal substantial non-encoded conformational variability within the Cas9 core, showing that this benchmark embeds a known structural signal within broader structural heterogeneity that methods must confront in practice. To separate these axes of variation, we used systematic deep classification to generate curated particle subsets depleted of selected domain motions while retaining the encoded labels. We further provide pooled particle stacks with standardized per-particle poses in a common reference frame and a lightweight framework for *in silico* particle pooling to generate challenge datasets with user-defined ground-truth distributions of encoded and non-encoded structural heterogeneity. Together, this resource supports robust benchmarking of heterogeneous reconstruction algorithms and provides a biochemically tractable model system for evaluating entire cryoEM pipelines, including alternative data-collection and preprocessing approaches, under experimentally realistic conditions.

## INTRODUCTION

High-resolution single-particle cryogenic electron microscopy (cryoEM) has become a routine route to determining near-atomic-resolution structures of macromolecular complexes (Callaway 2015). The same advances that enabled this transition, including direct electron detectors (Li *et al*. 2013), improved microscope stability (Fréchin *et al*. 2023) and automation (Mastronarde 2005; Suloway *et al*. 2005), and increasingly mature data processing pipelines (Bell *et al*. 2016; Punjani *et al*. 2017; Grant *et al*. 2018; Burt *et al*. 2024), have also shifted the field from producing a single consensus reconstruction toward resolving *ensembles* of conformational and compositional states (Elad *et al*. 2008; Frank 2009; Kinman *et al*. 2023; Webster *et al*. 2024). In practice, heterogeneity is now interrogated using a spectrum of approaches. These include discrete 2D- and 3D-classification (Scheres 2016), often using masks to focus on a specific region of the structure (Lyumkis *et al*. 2013; Nakane and Scheres 2021), and continuous models that map particles onto low-dimensional manifolds (Punjani and Fleet 2021; Gilles and Singer 2025), increasingly including deep-learning–based methods (Chen and Ludtke 2021; Zhong *et al*. 2021; Luo *et al*. 2023).

This rapid progress has, in turn, created a second-order challenge: heterogeneous reconstruction methods are difficult to evaluate and improve without benchmarks where the “right answer” (*i*.*e*., ground truth) is known. Many widely used benchmarks rely on datasets collected to analyze specific biological questions (Ru *et al*. 2015; Davis *et al*. 2016; Plaschka *et al*. 2017), where success is defined retrospectively based on whether a pipeline reproduces states recovered by alternative computational methods, or reveals a new state that appears biologically plausible. In favorable cases, such novel states are corroborated by orthogonal biochemical or genetic evidence (Sun *et al*. 2023). In many other cases, however, the benchmark becomes circular: a class is treated as “validated” because multiple pipelines recover it, even when its true frequency is unknown. Moreover, because per-particle labels are not independently available, such datasets are ill-suited for assessing per-particle classification accuracy.

Two trends make this limitation increasingly consequential. First, the field has moved from qualitative claims that alternative states exist to quantitative claims about their relative abundance, often supported by data from complementary assays (Fischer *et al*. 2010; May *et al*. 2026). These distributions are increasingly interpreted through the lens of kinetics and thermodynamics (Dashti *et al*. 2014; Giraldo-Barreto *et al*. 2021). Such inferences require not only sensitivity to the existence of a given state, but the accurate recovery of proportions of these states, and faithful estimates of how the proportions change across time or condition (Amann *et al*. 2023). As such, robust benchmarks then necessarily require ground-truth labels on a per-particle basis. Second, many modern approaches explicitly trade interpretability for representation power of the underlying model (Toader *et al*. 2023), for example by using deep-learning-based non-linear models (Zhong *et al*. 2021; Luo *et al*. 2023) instead of more interpretable linear models (Punjani and Fleet 2021; Gilles and Singer 2025). We expect that benchmarks with known labels will be critical in evaluating such tradeoffs across tools and laboratories.

Synthetic datasets address some of these issues by providing exact labels because one can simulate projections from known models, mix them in defined proportions, and compare reconstructions and particle assignments directly to ground truth. Such approaches have long existed (van Heel 1984; Sigworth 1998). Although these simulations are invaluable for rapidly testing algorithmic ideas, these purely synthetic benchmarks typically omit or simplify key nuisance parameters. For example, many simulators adopt a simple additive white Gaussian noise model (Baxter *et al*. 2009; Powell and Davis 2024; Jeon *et al*. 2024), because more physically realistic image-formation and noise simulations (Himes and Grigorieff 2021) remain computationally expensive. Further, in real datasets, orientations and defocus are imperfectly inferred from noisy images (Sigworth 1998; Mindell and Grigorieff 2003); the datasets contain contaminants and errantly picked particles (Voss *et al*. 2009); and 3D reconstructions often exhibit angular anisotropy (Tan *et al*. 2017) as a result of particles exhibiting a preferred orientation on the grid (Noble *et al*. 2018). Collectively, these experimental effects are difficult or impossible to model faithfully in synthetic benchmarks, yet they often dominate practical failure modes in analysis of structural heterogeneity (Evans *et al*. 2026).

Moreover, in real specimens, the interesting structural variability is typically embedded in a broader conformational ensemble rather than appearing in isolation (Frauenfelder *et al*. 1991; Henzler-Wildman and Kern 2007; Boehr *et al*. 2009). Indeed, multiple modes of conformational and compositional variability often coexist in these datasets; as a result, biologically uninteresting signals can often overwhelm subtler signals that may be of greater biological interest (Zhang *et al*. 2019). A benchmark that contains only a single encoded structural perturbation would therefore overestimate practical performance, whereas embedding a known signal within a background of non-encoded heterogeneity forces methods to separate the targeted changes from experimentally realistic confounding variables (*i*.*e*., nuisance parameters).

Motivated by these observations, we reasoned that a useful *experimental* benchmark dataset for analysis of structural heterogeneity should satisfy several criteria. It should include: (*i*) an encoded, labeled form of heterogeneity whose structural signature is known *a priori* and whose mixture proportions are controllable; (*ii*) realistic, *non-encoded* heterogeneity (*e*.*g*., conformational variability or partial occupancy) that methods must contend with in practice, ideally with options to deplete these confounding variables to titrate the difficulty of the task; (*iii*) realistic imaging artifacts and detector noise; (*iv*) nuisance-parameter uncertainty arising from realistic errors in estimation of the contrast transfer function (CTF) and of particle orientations; and (*v*) contaminants and imperfect particle picks that often consume modeling capacity and can limit performance. Finally, such a benchmark should be useful beyond algorithm comparisons: a well-designed biochemical sample should be able to serve as a test object for end-to-end pipeline evaluation, including data collection strategies, preprocessing choices (*e*.*g*., motion correction, CTF estimation, particle picking) as well as downstream refinement and heterogeneity analysis— effectively an “apoferritin-like” reference (Nakane *et al*. 2020; Yip *et al*. 2020; Zhang *et al*. 2020; Danev *et al*. 2021), but for heterogeneous specimens.

Here we developed such a benchmark using a catalytically inactive mutant of the *Streptococcus pyogenes* Cas9 nuclease (dCas9) bound to a guide RNA (sgRNA) and target DNA (tDNA) (Jinek *et al*. 2012). dCas9 is a biochemically tractable ~160 kDa multidomain protein that samples a rich conformational landscape, including domain-wide motions of the REC and HNH nuclease lobes (Jinek *et al*. 2014; Zhu *et al*. 2019). It thus recapitulates the kind of domain-scale heterogeneity that modern methods aim to resolve. At the same time, dCas9•sgRNA recognizes a fixed sequence (*i*.*e*., the protospacer/PAM core) on the tDNA while tolerating variation outside these recognition elements (Anders *et al*. 2014), creating a natural handle for engineering defined structural perturbations into the tDNA. We expected this architecture to mimic a realistic challenge in which classification and reconstruction methods must recover a localized structural signal while contending with non-encoded conformational variability within the same particle population, a setting likely well-matched to focused-classification strategies (von Loeffelholz *et al*. 2017).

Using this dCas9 system, we encoded ground-truth heterogeneity by engineering a series of tDNA duplexes with defined extensions, while holding the sgRNA and the protospacer/PAM-containing segment constant. Because each complex was assembled and vitrified independently, the dataset-of-origin provided an unambiguous label for every particle. We found that homogeneous refinements of individual datasets recovered the engineered tDNA extension signal, enabling the construction of a simple quantitative “tDNA extension occupancy” readout. These same reconstructions contained the non-encoded conformational variability within the dCas9 core we expected, which allowed us to curate particle subsets that depleted selected confounding variability while retaining encoded labels. Finally, we compiled pooled particle stacks with carefully refined pose estimates, and developed a lightweight framework for *in silico* particle pooling, which facilitates generating “challenge datasets” with user-defined ground-truth distributions. Together, we expect these resources will newly enable principled benchmarking of heterogeneity methods under realistic conditions.

## RESULTS

### Design and imaging of dCas9 complexes bearing encoded heterogeneity with biochemically defined per-particle labels

To realize this experimental benchmarking tool, we designed a series of dCas9•sgRNA•tDNA complexes in which the encoded perturbation was the length of an extension on the tDNA duplex (see Methods), while the sgRNA sequence and the protospacer/PAM-containing segment of the tDNA were held constant (**Figure 1A**). Our dCas9 construct also incorporated the inactivating D10A and H840A mutations (Jinek *et al*. 2012), which allow for tDNA binding but restrict cleavage (Qi *et al*. 2013). This design facilitated independent purification of sgRNA and dCas9 (**Figure 1B**), followed by dCas9•sgRNA•tDNA holocomplex assembly with a series of highly purified, commercially sourced tDNAs (see Methods). The resulting complexes were further purified by size-exclusion chromatography to remove excess tDNA (**Figure 1C**). Mass photometry (Asor *et al*. 2025) analysis of each complex identified a dominant, monodisperse species at the expected dCas9•sgRNA•tDNA molecular weight (**Figure 1D**).

**Figure 1.**
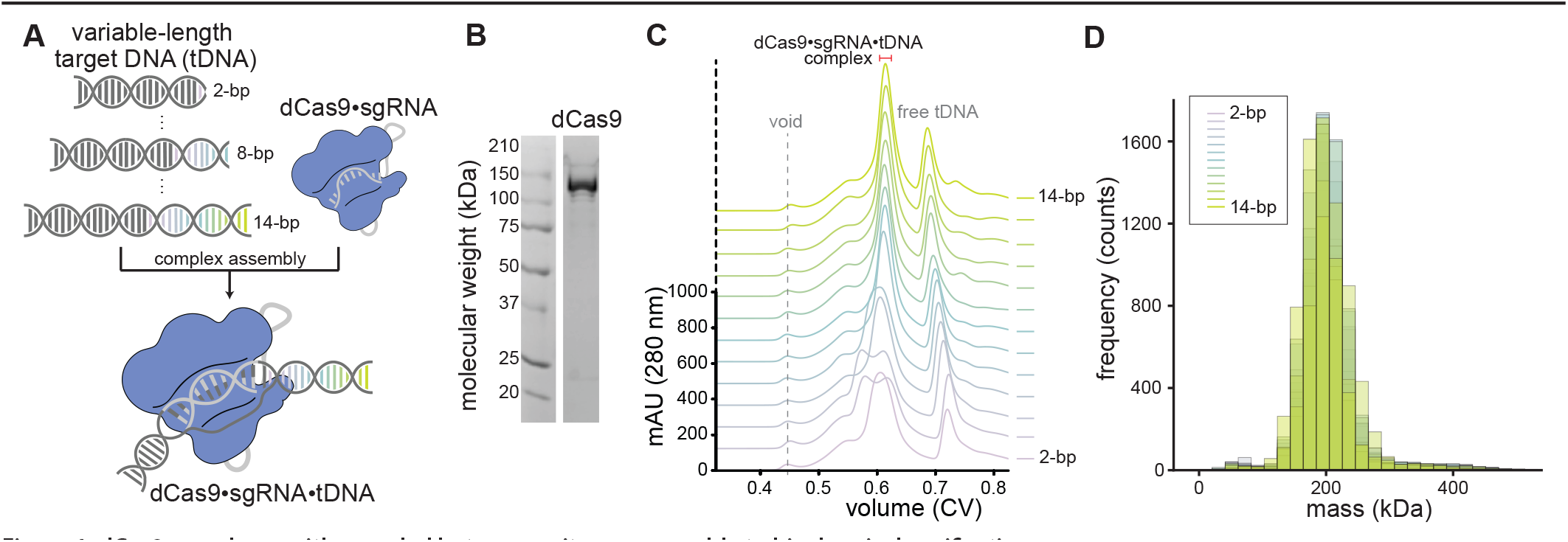
dCas9 complexes with encoded heterogeneity are amenable to biochemical purification. **(A)** Schematic illustrating the design of dCas9•sgRNA•tDNA complexes bearing variable-length tDNA extensions. **(B)** SDS-PAGE of purified dCas9, with molecular weight standards indicated. **(C)**Size-exclusion chromatography traces of dCas9•sgRNA•tDNA complexes monitored by absorbance at 280 nm. Traces are offset along the y-axis for clarity and colored according to the key at right. Major elution peaks are annotated; the red bar indicates fractions collected for mass photometry analysis and cryoEM grid preparation. **(D)** Mass distributions for each purified complex, as analyzed by mass photometry, shown as overlaid histograms of particle counts as a function of molecular mass. Histograms are colored according to the key. The expected mass of the holocomplex ranges 206–212 kDa for the 2-bp to 14-bp samples.

Each sample was next vitrified on a separate graphene-coated grid (Grassetti *et al*. 2023) and imaged under matched collection conditions (**Table S1)**, including +18° stage tilt that was employed to mitigate preferred orientation (Tan *et al*. 2017). Across the series, ~4,500–7,100 movies were collected per dataset, which we expected would provide sufficient particle counts for high-resolution refinement of individual datasets as well as pooled analyses at the scale of millions of particles. All datasets were pre-processed through a unified pipeline (**Supplementary Figure 1**; see Methods) to minimize dataset-specific optimization and ensure that differences observed across datasets primarily reflected the benchmark design as opposed to bespoke pre-processing choices. This approach resulted in ~10^5^-10^6^ particles per dataset (**Supplementary Figure 2**), which we expected would be sufficient for our planned analyses.

### Homogeneous refinement of individual datasets recovers the encoded tDNA extension signal

Independent homogeneous refinements in cryoSPARC yielded thirteen high-resolution maps (**Table S1**), with density for the tDNA extension qualitatively increasing across the series (**Figure 2A**). To quantify the tDNA extension signal, we masked this region and computed a normalized tDNA extension occupancy for each refined map (see Methods). Across the thirteen reconstructions, normalized occupancy largely increased monotonically with encoded tDNA extension length (**Figure 2B**). As such, we expect this tDNA occupancy metric could serve as a quantitative target for benchmarking. For example, in a dataset bearing a mixture of particle images with different tDNA lengths, a method that separates extension-length classes correctly should recover class maps with normalized tDNA extension occupancies matching those observed in each pure dataset.

**Figure 2.**
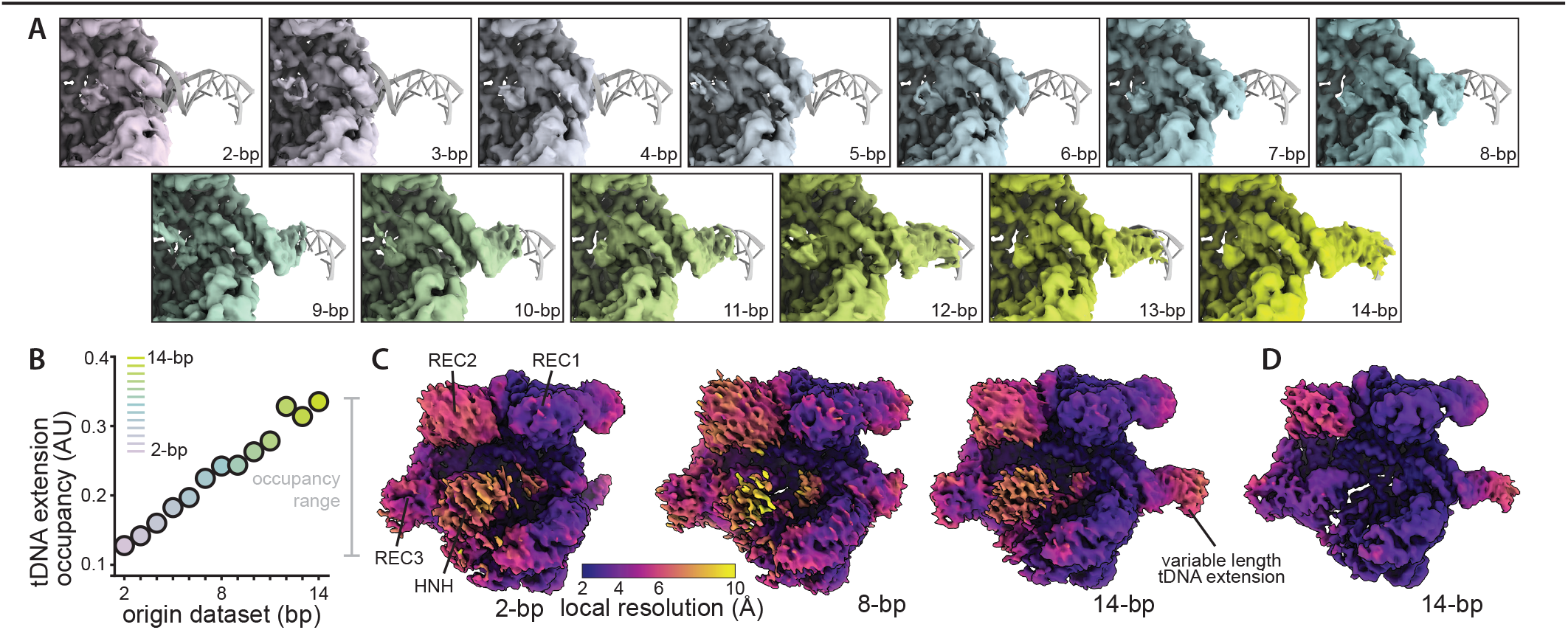
Benchmark datasets capture encoded and non-encoded heterogeneity. **(A)** Reconstructed density maps (surfaces) for each isolated dataset, colored by tDNA extension length from 2-bp (purple) to 14-bp (green), overlaid with the atomic model (gray ribbon) of the 14-bp dCas9•sgRNA•tDNA holocomplex. **(B)** Normalized tDNA extension occupancy for each reconstruction, plotted by the dataset of origin. Points are colored by dataset, following the key. The occupancy range, used here as a quantitative measure of the extent of observed heterogeneity, is indicated in gray. **(C)**Density maps colored by local resolution (Å) following key, shown at highly permissive contour levels to highlight reduced resolution in the conformationally variable REC2, REC3, and HNH domains. Representative reconstructions from the 2-bp (left), 8bp (middle), and 14-bp (right) datasets are shown. The variable-length tDNA extension is also indicated. **(D)** Density map reconstructed from the 14-bp particles exclusively, colored as in C, and displayed at a more stringent contour level to highlight the attenuated density of the HNH domain, which is consistent with large-scale conformational motions of this domain.

### Engineered dCas9 complexes retain conformational variability

In parallel, homogeneous refinements of these datasets exhibited the expected non-encoded heterogeneity within the dCas9 protein core, as evidenced by poorer local resolution estimates in the REC2, REC3, and HNH lobes (**Figure 2C**). Moreover, the HNH domain exhibited notably attenuated density relative to the other dCas9 domains and disappeared at more stringent contour levels (**Figure 2D**), likely a result of large conformational motions in this domain. These patterns persisted across extension lengths (**Supplementary Figure 3**), indicating that they are not a consequence of the encoded tDNA extensions. In practical terms, these results implied that even a single-label particle stack did not correspond to a “single structure”, but instead represented a structural ensemble, with non-encoded conformationally variable regions of the map overlaid on the set of fixed tDNA extensions.

To assemble target datasets for heterogeneous reconstruction, we next generated two large particle stacks bearing mixtures of tDNA extensions: a three-component mixture containing only the 2-bp, 8-bp, and 14-bp datasets (811,374 particles), and a full mixture containing particles from all thirteen datasets (2,904,834 particles) (**Table S2**). Homogeneous refinements of these mixed particle stacks produced high-resolution structures (2.5 Å GS-FSC and 2.3 Å GS-FSC, respectively; **Table S2**) and, as above, signs of conformational variability in the REC2, REC3, and HNH lobes were present (**Supplementary Figure 4**). Additionally, we observed attenuation of the density at the extremes of the tDNA extension region, as would be expected from averaging across the mixture of tDNA extension lengths.

### Deep hierarchical 3D-classification produces curated particlesubsetsdepletedofnon-encodedheterogeneity

Whereas particle stacks bearing complex structural mixtures better reflect the natural systems that heterogeneous classification and reconstruction tools aim to model, we recognized the value of simpler model systems whose structural heterogeneity was dominated by a single encoded source of variability. To build such a set, we performed deep classification of the pooled 2-14bp particle stack (2,904,834 particles) using a focused mask that excluded the tDNA extension region (see Methods). This approach yielded an ensemble of 512 volumes that captured conformational variation in dCas9 while holding the tDNA extension region effectively “invisible” to the classifier (**Figure 3A**).

**Figure 3.**
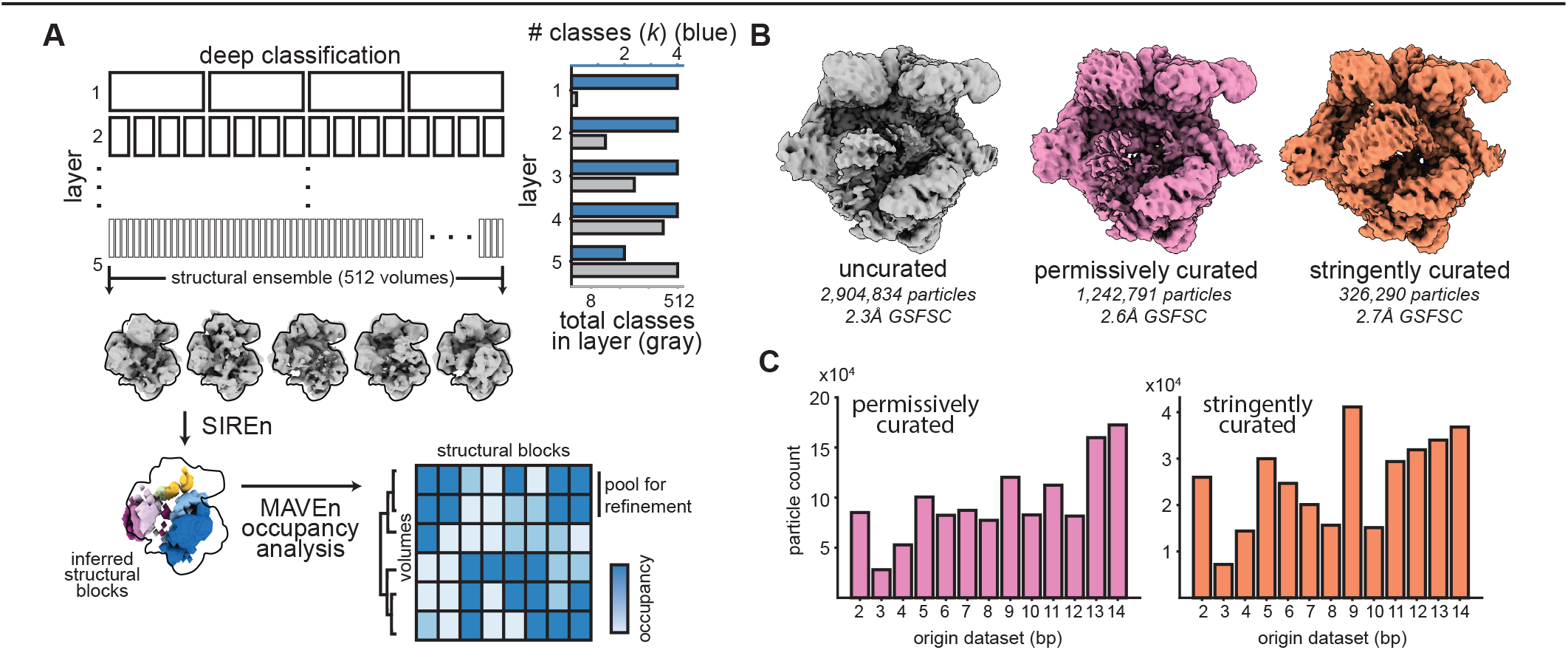
Curated subsets are depleted of non-encoded heterogeneity. **(A)** Deep classification, pooling, and refinement approach used to generate curated particle subsets depleted of non-encoded structural heterogeneity (see Methods). Representative classes from the volume ensemble produced by deep classification are shown in gray. Representative SIREn blocks used for querying occupancy are shown below, with schematic heatmap depicted to illustrate clustering approach. The real heatmap used to curate the permissive and stringent sets is shown in **Supplementary Figure 5**. **(B)** Refined maps of all 2-14 bp particles (left, gray), the permissively curated subset (middle, pink), and the stringently curated subset (right, orange). **(C)**Total particles from each origin dataset in the two curated subsets, colored as in (B).

To identify the dominant regions of structural variability in this ensemble, we applied SIREn (Kinman *et al*. 2025), which cleanly identified structural blocks corresponding to the disparate conformations of the REC2, REC3, and HNH domains (**Supplementary Figure 5A**). We then quantified the occupancy within each structural block across the 512 volumes using MAVEn (Sun *et al*. 2023), producing a matrix of normalized block occupancies that summarized the conformational state of each volume across the ensemble. Hierarchical clustering of this occupancy matrix produced 48 clusters (**Supplementary Figure 5B**), each corresponding to a set of structurally similar particles.

From these clusters, we curated two particle subsets that depleted non-encoded heterogeneity to different degrees, thereby producing classification tasks that one might expect to vary in difficulty. The “permissively curated” set, which included 1,242,791 particles, primarily decreased variability in the REC3-lobe, whereas the “stringently curated” particle stack aimed to select for a single dominant conformation for each of the HNH, REC2, and REC3 lobes (**Figure 3B**). This more stringent curation resulted in 326,290 particles. Homogeneous refinement and local resolution analyses of these curated subsets were consistent with this selection strategy having achieved its aims - the curated maps successively exhibited less density attenuation across the REC and HNH lobes (**Supplementary Figure 6**), all while maintaining variance in the tDNA extension region as assessed by the number of particles contributed by each dataset (**Figure 3C**). An additional practical output of the deep-classification pipeline was refined pose estimates for each particle. Per-particle pose estimates were largely consistent across the two approaches (**Supplementary Figure 7A)**, but for a small subset of particles, the poses estimated by deep classification differed significantly from those estimated in the consensus refinement (**Supplementary Figure 7B**).

#### *In silico* mixing scripts enable the generation of controlled challenge datasets

Having curated these particle subsets, we recognized the value of generating particle mixtures designed to test specific aspects of heterogeneous structure analysis (*e*.*g*., highly skewed tDNA extension distributions). To this end, we developed a lightweight script that resamples the full set of all particles across each of the 13 datasets. As each particle is labeled by dataset-of-origin and inferred conformational state from deep classification and pooled refinement, this script allows users to produce new mixed stacks according to user-defined distributions of encoded and non-encoded heterogeneity (**Figure 4**). This tool enables systematic construction of challenges such as: balanced mixtures spanning many extension lengths; skewed mixtures in which a particular extension length is rare; and mixtures that couple or decouple the encoded extension label from dominant non-encoded conformational classes. Taken together, we expect that these datasets and our *in silico* mixing tools will enable systematic construction of challenge datasets that isolate specific failure modes, providing a controlled setting for assessing and comparing heterogeneous reconstruction methods.

**Figure 4.**
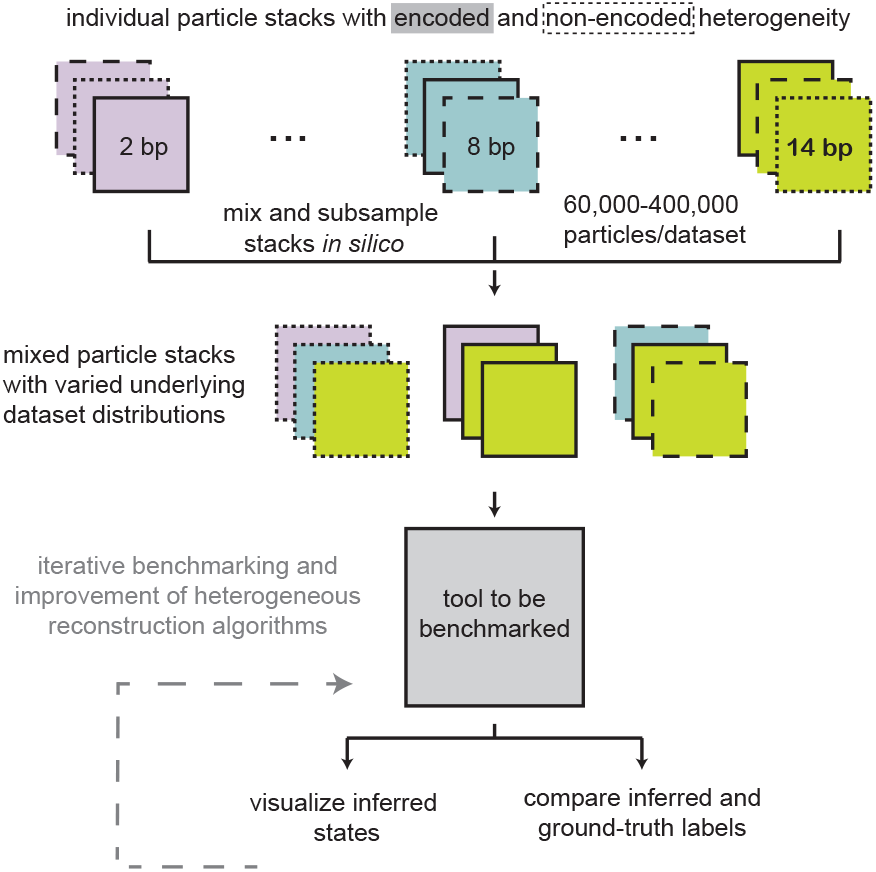
Cas9 datasets can be subsampled and mixed *in silico* to challenge heterogeneous reconstruction algorithms. Schematic illustrating the approach implemented in `generate_ distributions.py script` to create mixed particle stacks with varied underlying dataset distributions.

## DISCUSSION

The goal of this work is to provide an experimentally grounded benchmark for heterogeneous cryoEM analysis that preserves both a known structural signal and the confounding variability intrinsic to real datasets. By encoding heterogeneity through a defined tDNA extension while retaining non-encoded conformational variability within dCas9, this system enables evaluation of method performance under conditions that more closely reflect practical use. The resulting datasets provide unambiguous labels via dataset-of-origin that can be used to assess particle classification accuracy, while the extension-occupancy metric provides a simple, quantitative readout for reconstruction fidelity in the tDNA extension. Critically, this framing separates the problem of detecting and quantifying heterogeneity from that of validating its biological interpretation.

We expect that the ease of preparing these dCas9•sgRNA•tDNA complexes will additionally make them well suited for evaluating experimental choices under controlled conditions. For example, additional grids could be prepared and imaged using a set of alternative data-collection strategies while keeping the downstream analysis unchanged. Emblematic of such an approach, we reimaged a subset of these samples with an alternative microscope and detector (**Table S4**) and we are making these datasets publicly available for analysis. In this setting, differences in performance could be directly attributed to the data-collection strategies, as measured by the aforementioned metrics of accuracy in per-particle classification and recovery of the encoded tDNA structure. This approach would additionally enable systematic comparison of more dramatically altered data-collection strategies in a way that is not possible in typical biological datasets, where the ground truth is unknown and experimental and computational effects are intertwined.

Beyond benchmarking reconstruction accuracy, these datasets also provide a controlled setting to assess the effects of pose estimation on heterogeneous reconstruction algorithms. By supplying standardized per-particle poses derived from a consistent classification and refinement pipeline, the benchmark supports comparisons in which alignment is held fixed, thereby isolating the performance of downstream heterogeneity analysis. Conversely, because raw data and intermediate processing steps are available, the same system can be used to evaluate end-to-end workflows in which pose estimation and classification are jointly optimized. In both settings, improvements can be quantified through their recovery of the encoded signal, providing a concrete link between alignment accuracy and downstream interpretability.

The inclusion of curated particle subsets further enables mechanistic interrogation of method performance. By selectively depleting dominant sources of non-encoded conformational variability, these subsets create a ladder of difficulty that we hope developers can use to distinguish between methodological limitations in sensitivity to the encoded signal versus methodological susceptibility to confounding heterogeneity. In this framework, progress can be measured as methods advance from accurate classification of the curated subsets toward correct recovery of structure in the full, non-curated datasets.

More broadly, this work suggests a path toward a richer ecosystem of experimentally grounded datasets with known ground truth. One direction is to expand the repertoire of encoded perturbations within this dCas9 framework - for example, by engineering domain deletions or targeting multiple dCas9 molecules to variable-length tDNAs with multiple binding sites. Another is to develop analogous systems using different programmable protein or RNA scaffolds, including those suitable for *in situ* cryo-electron tomography, where ground-truth benchmarks are largely absent. Taken together, these datasets shift benchmarking from retrospective validation of plausible states to prospective evaluation against a known, experimentally encoded signal embedded within realistic structural variability, uncertainty, and noise. In doing so, they aspire to provide a foundation for the development of more quantitative and reproducible methods for resolving structural heterogeneity in cryoEM.

## DATA AVAILABILITY

Raw movies and particle stacks for all described datasets will be deposited at EMPIAR upon publication. Density maps for individual 2-bp – 14-bp refinements and curated subsets have been deposited at EMDB. MAVEn and SIREn results are available at zenodo.org/records/19102039.

## CODE AVAILABILITY

Scripts used for subsampling particle images are available at https://github.com/lkinman/benchmark_datasets.

## ACKNOWLEDGEMENTS

This research was supported by the NIH grant R01GM144542, NSF-CAREER grant 2046778, and the Smith Family Odyssey Award (JHD). Samples were prepared at the MIT.nano cryoEM facility and screened on a Talos Arctica microscope, which was a gift from the Arnold and Mabel Beckman Foundation. A portion of this research was supported by NIH grant R24GM154185 and performed at the Pacific Northwest Center for Cryo-EM (PNCC) with assistance from Vamseedhar Rayaprolu. Molecular graphics and analyses performed with UCSF ChimeraX, developed by the Resource for Biocomputing, Visualization, and Informatics at the University of California, San Francisco, with support from NIH R01GM129325 and the Office of Cyber Infrastructure and Computational Biology, NIAID. The authors used AI-assisted language tools (ChatGPT, OpenAI) for grammatical and stylistic copy-editing; all scientific content was written, verified, and is the sole responsibility of the authors.

## AUTHOR CONTRIBUTIONS

Conceptualization: All authors.

Data curation: LFK, AVG.

Methodology: AVG.

Investigation: AVG, LFK.

Software: LFK.

Supervision: JHD.

Visualization: All authors.

Writing – original draft: JHD.

Writing – review & editing: All authors.

Funding acquisition and project administration: JHD.

## MATERIALS AND METHODS

### Construct design

A plasmid expressing catalytically inactive dCas9 (10xHis-MBP-TEV-S. pyogenes dCas9 M1C D10A C80S H840A C574S) was a gift from Jennifer Doudna (Addgene plasmid #60815). sgRNA was purchased from Synthego with the sequence:

5’-GACGCAUAAAGAUGAGACGCGUUUUAGAGCUAGAAAUAGCAAGUUAAAAUA-AGGCUAGUCCGUUAUCAACUUGAAAAAGUGGCACCGAGUCGGUGCUU-3’

Target DNA duplexes were ordered as PAGE-purified oligos from IDT; sequences are provided in **Table S3**.

### dCas9 purification

The Cas9 plasmid was transformed into the *E. coli* strain Rosetta 2 (Novagen) for protein expression. Rosetta 2 cells were grown in 2xYT media supplemented with 25 mg/mL chloramphenicol and 100 mg/mL ampicillin at 37 °C until they reached an optical density (600 nm) of 0.6, at which point Isopropyl β-D-1-thiogalactopyranoside (IPTG) was added to a final concentration of 200 mM and the temperature was reduced to 16 °C for 16 hours. Cell pellets were collected by centrifugation at 4,000 x g for 15 minutes before snap freezing in liquid nitrogen and storage at −80 °C. Cell pellets were resuspended in buffer LB (20 mM HEPES pH 7.5, 500 mM NaCl, 5 mM 2-mercaptoethanol), supplemented with EDTA-free protease inhibitor tablets) and lysed using a cryo-mill (Retsch). Cell lysates were then sonicated (Branson SFX250) with a ¼” tip at 25% amplitude with 10-second-on / 20-second-off cycles for a total of 2 minutes. Lysate was clarified by centrifugation at 38,000 x g for 60 minutes. Clarified lysate was incubated for 30 minutes at 4 °C with 1.5 mL Ni Sepharose 6 Fast Flow (GE 17-5318-02) resin per 1 L of cell culture. Resin was allowed to settle in a gravity column (BioRad), washed with 25 column volumes (CV) of lysis buffer, and eluted in buffer EB (Tris pH 8.1, 250 mM NaCl, 5 mM BME, 10% glycerol and 350 mM imidazole). To remove the 10xHis-MBP tag, dCas9 protein was dialyzed overnight at 4°C into buffer DB (20 mM HEPES pH 7.5, 100 mM KCl, 5 mM BME, 10% glycerol) in the presence of recombinant TEV-protease (S219V) at a molar ratio of 2.5:1 dCas9:TEV. Dialyzed dCas9 was further purified using a 5 mL HiTrap SP FF strong cation exchange column (Cytiva 17515701). dCas9 elution fractions from SP were combined and concentrated at 4000 x g with a VIVASPIN-20 50 kDa MW cutoff spin concentrator (Sartorius #VS2032). Concentrated protein was applied to a HiLoad 16/600 Superdex 200 pg (Cytiva 28989335) column in buffer GF (20 mM HEPES pH 7.5, 100 mM KCl, 5 mM BME, 10% glycerol).

### Complexation and biochemical validation

DNA target oligos were mixed at equimolar ratios in buffer HB (20 mM HEPES pH 7.5, 100 mM KCl, 5 mM MgCl2, 3% glycerol), heated at 95 °C for 2 min, then slowly cooled on benchtop. Catalytically inactivated Cas9 (dCas9) and sgRNA were mixed at a 1:1 molar ratio (25 µM) in buffer CB (20 mM HEPES pH 7.5, 100 mM KCl, 5 mM MgCl_2_, 1 mM DTT) and incubated at 37 °C for 10 minutes. Samples were cooled to room temperature and DNA was added at a concentration of 50 µM. Samples were incubated for an additional 10 minutes at 37 °C. Cas9•sgRNA•DNA complexes were purified from unreacted components using an Agilent 1100 HPLC equipped with a Bio-SEC 3 (Agilent 5190-2513) size exclusion chromatography column run in buffer VB (20 mM HEPES pH 7.5, 100 mM KCl, 100 mM NaCl, 5 mM MgCl_2_, 1 mM DTT). Selected fractions were then analyzed by mass photometry. Briefly, a Refeyn mass photometer was calibrated with Novex NativeMark protein standards. Cas9•sgRNA•DNA samples were diluted five-fold to approximately 120 nM in buffer VB immediately prior to measurement by adding 3 µL of sample to 12 µL of buffer pre-loaded into analysis wells. Data from 60-120 second recordings were collected and analyzed using AcquireMP and DiscoverMP software.

### CryoEM sample preparation

Cas9-sgRNA-DNA complexes confirmed by mass photometry were applied to Quantifoil 2/1 Cu 200 grids at 125 ng/µL (~600 nM). The grids were coated with monolayer graphene, prepared as described previously (Grassetti *et al*. 2023), and plunged using a Vitrobot Mark IV (Thermo) vitrification system. Prior to plunging, grids were treated with UV/ozone for 7-8 minutes using a Bioforce PC440 UV/ozone cleaner. Vitrobot temperature was set to 10 °C and 95% humidity. Blotting force was set 9 units above the contact point of blotting pads without Whatman paper attached. Sample was incubated on grid for 10-20 seconds before blotting for 4 seconds and plunging into liquid ethane.

### CryoEM data collection

Datasets were collected at the Pacific Northwest Center for CryoEM (PNCC). Movies were collected on a Titan Krios equipped with a Falcon4i detector at 120,000x magnification without AFIS and with +18.0° stage tilt. Data collection parameters are outlined in **Table S1**. Data was stored in EER format. Additional datasets collected at MIT.nano were collected on a Titan Krios G3i at 130,000x magnification with AFIS and with +18.0° tilt. Data collection parameters for these datasets are outlined in **Table S4**.

### Initial preprocessing and particle picking with cryoSPARC

The final thirteen collected cryoEM datasets collected at PNCC were processed as outlined in **Supplementary Figure 1**. Briefly, raw movies were imported in cryoSPARC v4 (Punjani *et al*. 2017) for patch motion correction and patch CTF estimation. CryoSPARC’s blob picker tool was used to pick particles with a diameter of 115–135 Å. Particles were extracted with a box size of 896 pix (0.32 Å/pix) and Fourier cropped to 256 pix (1.12 Å /pix). *Ab initio* reconstruction was performed using all extracted particles and 3 total classes. Particles from the best class, as assessed by visual inspection, were selected for further processing.

### Final preprocessing, refinement, and polishing

Raw movies from each dataset were separately imported into RELION-5 (Burt *et al*. 2024) for preprocessing, particle extraction, and particle filtration. Motion correction and dose-weighting were performed with RELION’s own implementations of these tools; CTFFIND4 (Rohou and Grigorieff 2015) was used for CTF estimation. Filtered particle picks from multi-class *ab initio* reconstruction in cryoSPARC (described above) were re-extracted from micrographs in RELION following particle coordinate conversion with the csparc2star.py and star.py functions implemented in pyem (Asarnow *et al*. 2019). Particles were extracted at a box size of 448 pix (0.64 Å/pix) and Fourier cropped to 256 pix (1.12 Å/ pix). Re-extracted particles were subjected to one additional round of filtration by 2D classification before datasets were pooled for refinement. Two pooled refinements were done in RELION: one with particles from the 2-, 8-, and 14-bp datasets only, and one with particles from all thirteen datasets. Both refinements were done with the same parameters, using Blush regularization (Kimanius *et al*. 2024) and a reference volume that was initially low-pass filtered to 60 Å resolution. After initial refinement, particles were further refined, correcting first for higher-order aberrations, then anisotropic magnification, and finally per-particle defocus. Each particle stack was refined again, using the same parameters as before. Finally, each particle stack was polished (Zivanov *et al*. 2019) to correct for beam-induced motion and refined once more. Particle stacks were then imported into cryoSPARC for filtering by per-particle scale factor.

### Individual dataset refinements and scale-filtering

The polished particle stack containing particle images from all thirteen datasets (2-14 bp) was again subdivided to generate a separate stack for each individual dataset. The resulting thirteen stacks were imported into cryoSPARC v4.5.3. and subjected to non-uniform refinement. The distribution of per-particle scale factors from each refinement was fit using a two-component Gaussian mixture model (GMM), and labels (low-scale or high-scale) were predicted for each particle based on this fit GMM. Low-scale particles were discarded, while high-scale particles were retained, refined, and re-pooled for deep classification. The re-pooled particle stack was also imported into RELION for a final consensus refinement (**Figure 2, Supplementary Figure 4B**). The low-scale particles were also discarded from the 2/8/14-bp jointly-polished stack, and the resulting filtered stack was similarly reimported into RELION for a final consensus refinement (**Supplementary Figure 4A**).

### Deep classification and pooled refinement

Pooled datasets were subjected to systematic deep classification: 4 layers of classification with *k* = 4 classes per layer, followed by a single layer with *k* = 2 (producing 512 total classes in the final layer). To ensure particles sorted on non-encoded heterogeneity rather than the length of the DNA extension, a focus mask was used, covering the entire volume except the DNA extension. A filter resolution of 8 Å was used for all classification layers; parameters were otherwise kept at default values. SIREn (Kinman *et al*. 2025) was then used to identify regions within the volume that varied across the ensemble generated by deep classification. Each volume in the ensemble was then binarized using SIREn and queried for occupancy of each of the SIREn blocks with MAVEn (Davis *et al*. 2016; Davis and Williamson 2017; Kinman *et al*. 2023). Resulting occupancies were normalized by the total number of voxels within the SIREn block. Hierarchical clustering of volumes based on their occupancies for selected SIREn blocks partitioned the 512 classification volumes, and their associated particles, into 48 classes. All particles in a given class were pooled for refinement. To ensure that all refinements remained in the same reference frame, the final RELION consensus refinement from the 2/8/14-bp mixed particle stack, low-pass filtered to 30 Å `(common_frame_map`, hereafter), was used as the reference volume for all refinements. The final deposited particle stacks and .star files contain high-scale particles from the polished particle stacks (2/8/14-bp and 2-14 bp) with poses estimated by deep classification and pooled refinement approach. To quantitatively compare poses from the consensus refinement to poses from the deep classification and pooled refinement approach, we converted the φ and θ Euler angles into a 3D vector representing the projection angle of each particle on a unit sphere. For each particle, the angle α between these two vectors was calculated as: 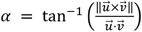

All particles with angular difference between 45° and 135° were designated as “high difference”.

### Generation of depleted subsets

Depleted subsets were generated by pooling particles from the 48 refined classes. To generate a “stringently curated” subset, in which the REC3, REC2, and HNH domains were conformationally homogeneous, particles from classes 26-30 were pooled and subsequently refined in cryoSPARC, using the `common_frame_map as` an initial model. To generate a “permissively curated” subset, in which only REC3 heterogeneity was depleted, classes 0-25 were pooled and refined as above (**Supplementary Figures 5-6**).

### Atomic model building

An atomic model of the 14-bp extension was constructed following refinement of the 14-bp complex from a dataset collected at MIT.nano (**Table S4**). Briefly, the raw movies were imported into cryoSPARC (v4) for patch motion correction and patch CTF estimation. Initial particle picks were generated using the blob picker tool, with particle diameter between 100 and 130Å. The resulting picks were extracted at a box size of 432 pix (0.65 Å/pix) and Fourier cropped to a box size of 128 pix (2.19 Å/pix). Selected class images after two rounds of 3D classification were used as templates for template picking. Particles from template picking were again extracted at a box size of 432 pix (0.65 Å/pix) and Fourier cropped to a box size of 128 pix (2.19 Å/pix). The resulting particle stack was filtered via 2D classification, re-extracted at a box size of 256 pix (1.10 Å/pix), filtered once more with 2D classification, and used to generate an initial model with a one-class *ab initio* reconstruction. The resulting model was refined using non-uniform refinement, and subsequently subject to unmasked 3D classification with 4 classes. Non-uniform refinement was applied one final time to the “best” class, identified by visual inspection, to generate the final map. To generate the atomic model, a previously published structure (Zhu *et al*. 2019) (PDB 6o0Z) was fit into the cryoEM density map as a rigid body using UCSF ChimeraX (Meng *et al*. 2023). DNA extensions were incorporated directly into this model using the Model/ Fit/Refine Add Terminal residue feature in Coot (Casanal *et al*. 2020). Refinement was carried out iteratively using ISOLDE (Croll 2018), through its ChimeraX plugin and Coot. ISOLDE was used for interactive fitting of the model into the density with molecular dynamics–based restraints. Coot was used to inspect the structure residue-by-residue to ensure that backbone placement and sidechain rotamers were consistent with the density. Side chains were modeled only where supported by clear density. If cryoEM density was insufficient to support a specific assignment, side chains were truncated, or omitted entirely.

## Supplementary Information

### SUPPLEMENTARY TABLES AND FIGURES

**Table S1.** CryoEM data collection and individual refinement parameters. Global resolution is provided at the 0.143 GS-FSC threshold. SCF* was calculated as described in Baldwin and Lyumkis, 2021.

| Dataset (bp) | 2 | 3 | 4 | 5 | 6 | 7 | 8 | 9 | 10 | 11 | 12 | 13 | 14 |
| --- | --- | --- | --- | --- | --- | --- | --- | --- | --- | --- | --- | --- | --- |
| Microscope | Titan Krios (PNCC) |  |  |  |  |  |  |  |  |  |  |  |  |
| Camera | Falcon 4i |  |  |  |  |  |  |  |  |  |  |  |  |
| Voltage (keV) | 300 |  |  |  |  |  |  |  |  |  |  |  |  |
| Magnification | 120,000x |  |  |  |  |  |  |  |  |  |  |  |  |
| Pixel size (Å) | 0.641 |  |  |  |  |  |  |  |  |  |  |  |  |
| Total dose (e<sup>-</sup>/Å<sup>2</sup>) | 55 | 55 | 55 | 55 | 55 | 55 | 55 | 55 | 55 | 55 | 55 | 55 | 55 |
| Dose rate (e<sup>-</sup>/pix/s) | 6.17 | 5.38 | 5.38 | 5.38 | 6.17 | 6.17 | 6.17 | 5.38 | 6.17 | 6.17 | 6.17 | 6.17 | 6.17 |
| Exposure time (s) | 3.89 | 4.45 | 4.45 | 4.45 | 3.89 | 3.89 | 3.89 | 4.45 | 3.89 | 3.89 | 3.89 | 3.89 | 3.89 |
| Defocus (µm) | -0.4, -0.6, -0.8, -1, -1.2, -1.4, -1.6, -1.8, -2.0, -2.2, -2.4 |  |  |  |  |  |  |  |  |  |  |  |  |
| Stage tilt (°) | +18.0 |  |  |  |  |  |  |  |  |  |  |  |  |
| Fractions | EER format |  |  |  |  |  |  |  |  |  |  |  |  |
| Movies (no.) | 7,100 | 4,563 | 5,836 | 4,682 | 6,073 | 5,357 | 5,137 | 5,308 | 6,048 | 5,019 | 4,949 | 5,474 | 5,507 |
| EMPIAR ID | Available upon publication |  |  |  |  |  |  |  |  |  |  |  |  |
| Final particles (no.) | 215,016 | 63,528 | 130,527 | 224,116 | 221,412 | 222,120 | 197,780 | 261,551 | 173,173 | 259,355 | 179,796 | 357,882 | 398,578 |
| Final pixel size (Å) | 1.12 |  |  |  |  |  |  |  |  |  |  |  |  |
| EMDB ID | 76407 | 76406 | 76405 | 76404 | 76403 | 76402 | 76401 | 76400 | 76399 | 76398 | 76397 | 76396 | 76395 |
| Global resolution | 2.98 | 3.09 | 3.08 | 2.98 | 2.97 | 3.06 | 2.97 | 2.80 | 2.98 | 2.91 | 2.97 | 2.76 | 2.90 |
| Sharpening B-factor (Å<sup>2</sup>) | -83.6 | -61.8 | -79.5 | -85.0 | -82.5 | -88.7 | -79.1 | -78.5 | -86.2 | -80.3 | -70.5 | -80.9 | -95.7 |
| SCF* | 0.63 | 0.79 | 0.66 | 0.58 | 0.56 | 0.44 | 0.55 | 0.77 | 0.76 | 0.60 | 0.48 | 0.47 | 0.57 |

**Table S2.**
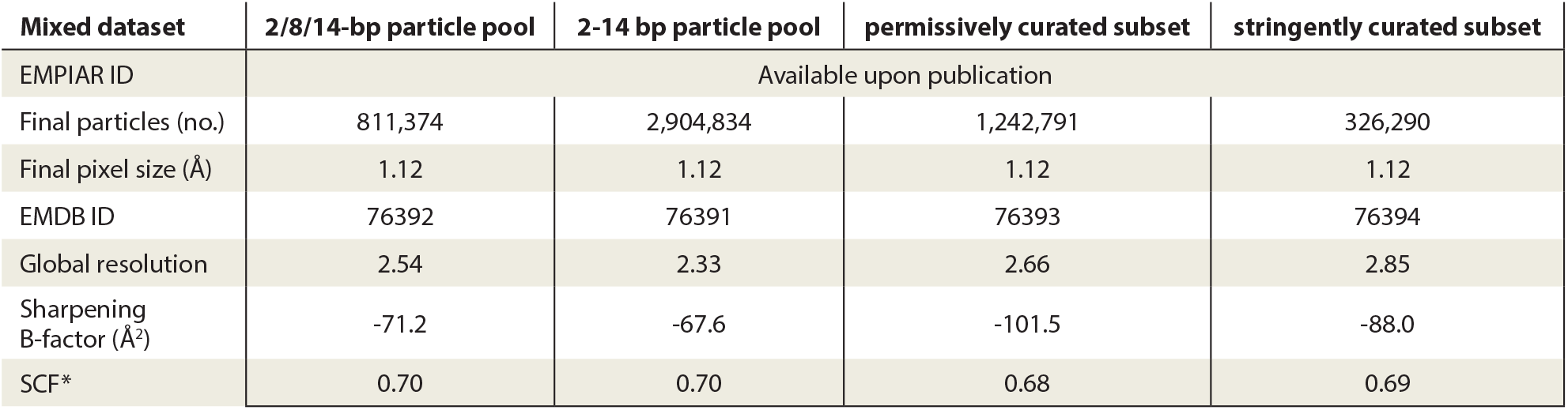
Deposited cryoEM particle stacks. Described particle stacks were deposited to EMPIAR, alongside raw movies (**Table S1**) and corresponding refinement metadata (*e*.*g*., pose, CTF parameters). Global resolution is provided at the 0.143 GS-FSC threshold. SCF* was calculated as described in Baldwin and Lyumkis, 2021.

| Mixed dataset | 2/8/14-bp particle pool | 2-14 bp particle pool | permissively curated subset | stringently curated subset |
| --- | --- | --- | --- | --- |
| EMPIAR ID | Available upon publication |  |  |  |
| Final particles (no.) | 811,374 | 2,904,834 | 1,242,791 | 326,290 |
| Final pixel size (Å) | 1.12 | 1.12 | 1.12 | 1.12 |
| EMDB ID | 76392 | 76391 | 76393 | 76394 |
| Global resolution | 2.54 | 2.33 | 2.66 | 2.85 |
| Sharpening B-factor (Å<sup>2</sup>) | -71.2 | -67.6 | -101.5 | -88.0 |
| SCF* | 0.70 | 0.70 | 0.68 | 0.69 |

**Table S3.**
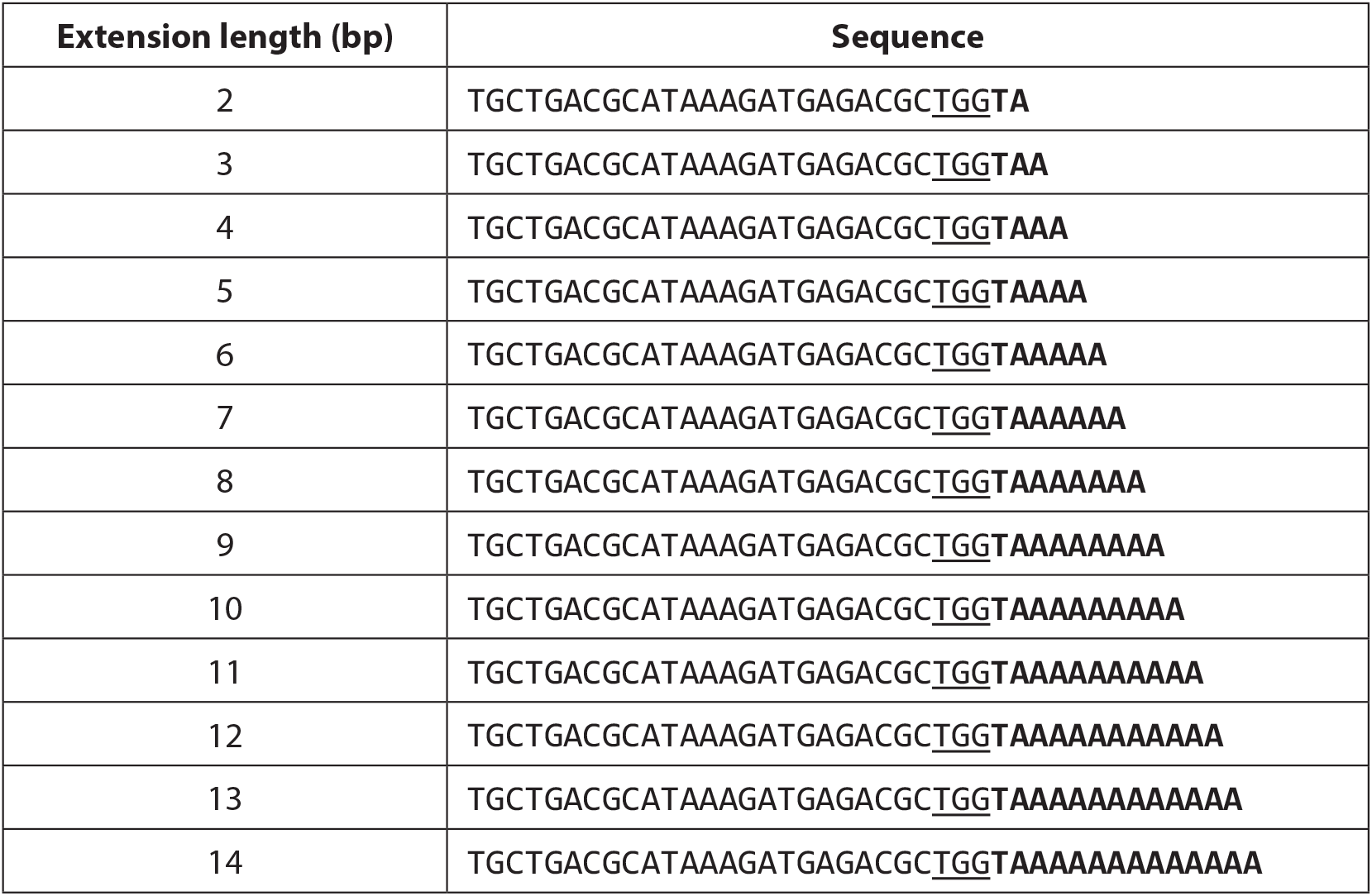
Nucleotide sequences for variable target DNA duplexes. Sequences are shown for only the forward strand. The protospacer adjacent motif (PAM) is underlined, and the tDNA extension is in bold.

**Table S4.** CryoEM data collection parameters using alternative instrumentation.

| Dataset (bp) | 2 | 3 | 4 | 5 | 6 | 7 | 8 | 14 |
| --- | --- | --- | --- | --- | --- | --- | --- | --- |
| Microscope | Titan Krios G3i (MIT.nano) |  |  |  |  |  |  |  |
| Camera | Bioquantum K3 |  |  |  |  |  |  |  |
| Voltage (keV) | 300 |  |  |  |  |  |  |  |
| Magnification | 130,000x |  |  |  |  |  |  |  |
| Pixel size (Å) | 0.654 |  |  |  |  |  |  |  |
| Total dose (e<sup>-</sup>/Å<sup>2</sup>) | 54.17 |  |  |  |  |  |  |  |
| Dose rate (e<sup>-</sup>/pix/s) | 21.7 |  |  |  |  |  |  |  |
| Exposure time (s) | 1.1 |  |  |  |  |  |  |  |
| Defocus (μm) | -0.4, -0.6, -0.8, -1, -1.2, -1.4, -1.6, -1.8, -2.0, -2.2, -2.4 |  |  |  |  |  |  |  |
| Stage tilt (°) | +18.0 |  |  |  |  |  |  |  |
| Fractions | 40 |  |  |  |  |  |  |  |
| Movies collected (no.) | 5,458 | 5,343 | 5,504 | 6,567 | 5,505 | 5,354 | 5,040 | 5,081 |
| EMPIAR ID | Available upon publication |  |  |  |  |  |  |  |

**Supplementary Figure 1.**
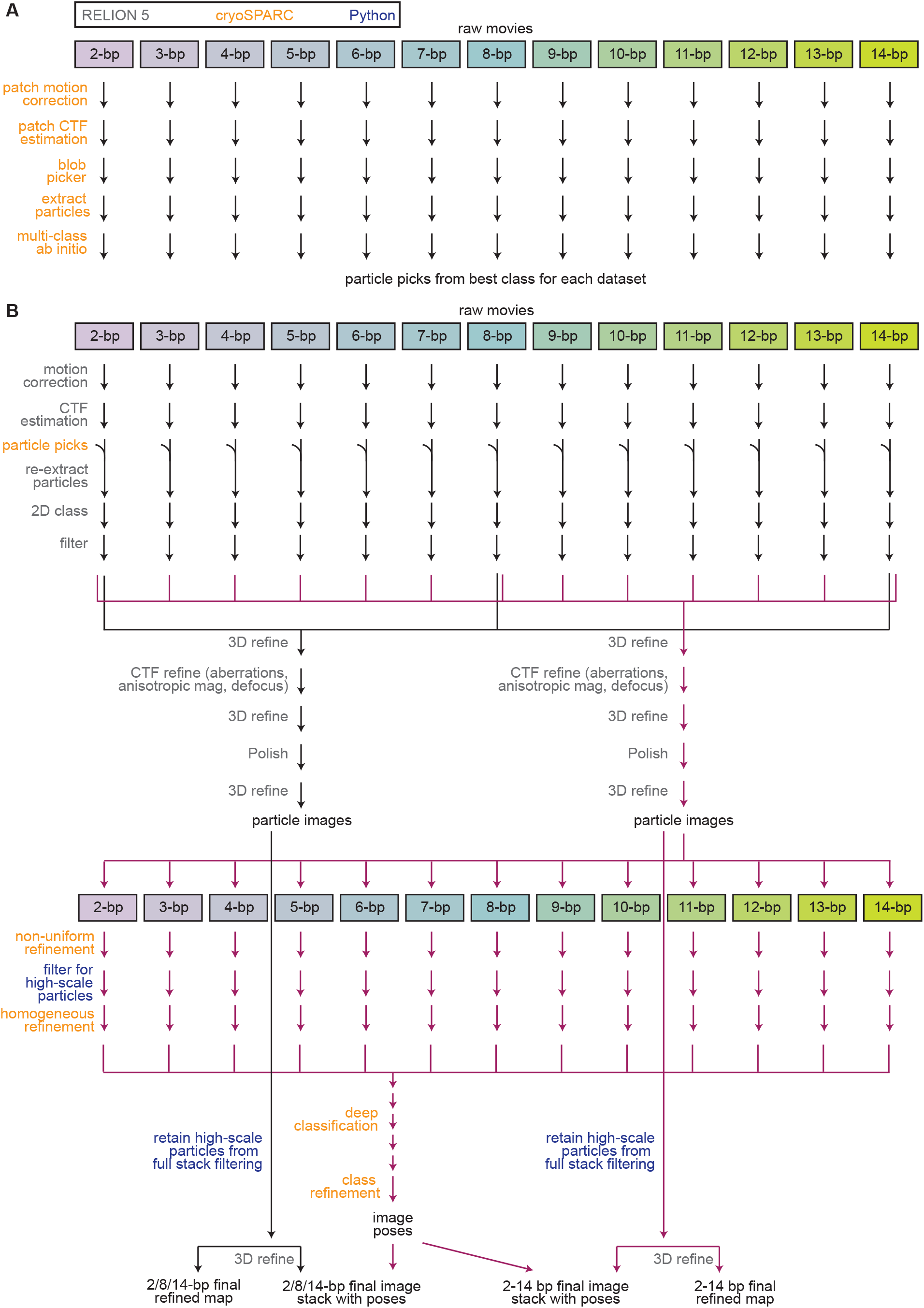
CryoEM data pre-processing approach. **(A)** Processing approach to generate initial particle picks in cryoSPARC. **(B)** Processing approach to generate final mixed particle stacks with accompanying poses from deep classification, alongside final refined consensus maps. Initial particle picks in (B) were generated as shown in (A) and imported into RELION using pyem (see Methods).

**Supplementary Figure 2.**
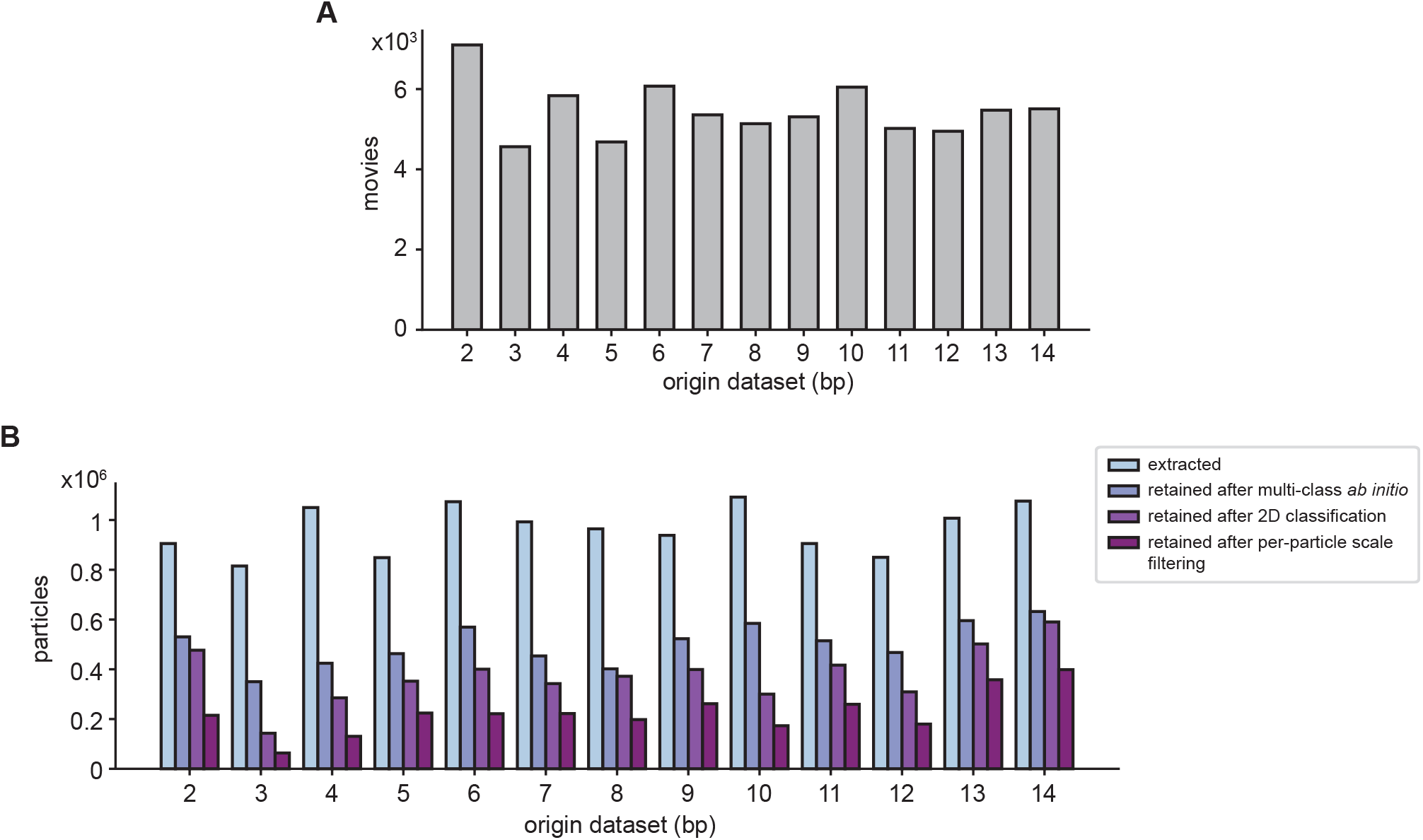
Per-dataset movie and particle statistics. **(A)** Number of raw movies collected for each dataset. **(B)** Number of particles originally picked and retained after each stage of filtering for each dataset.

**Supplementary Figure 3.**
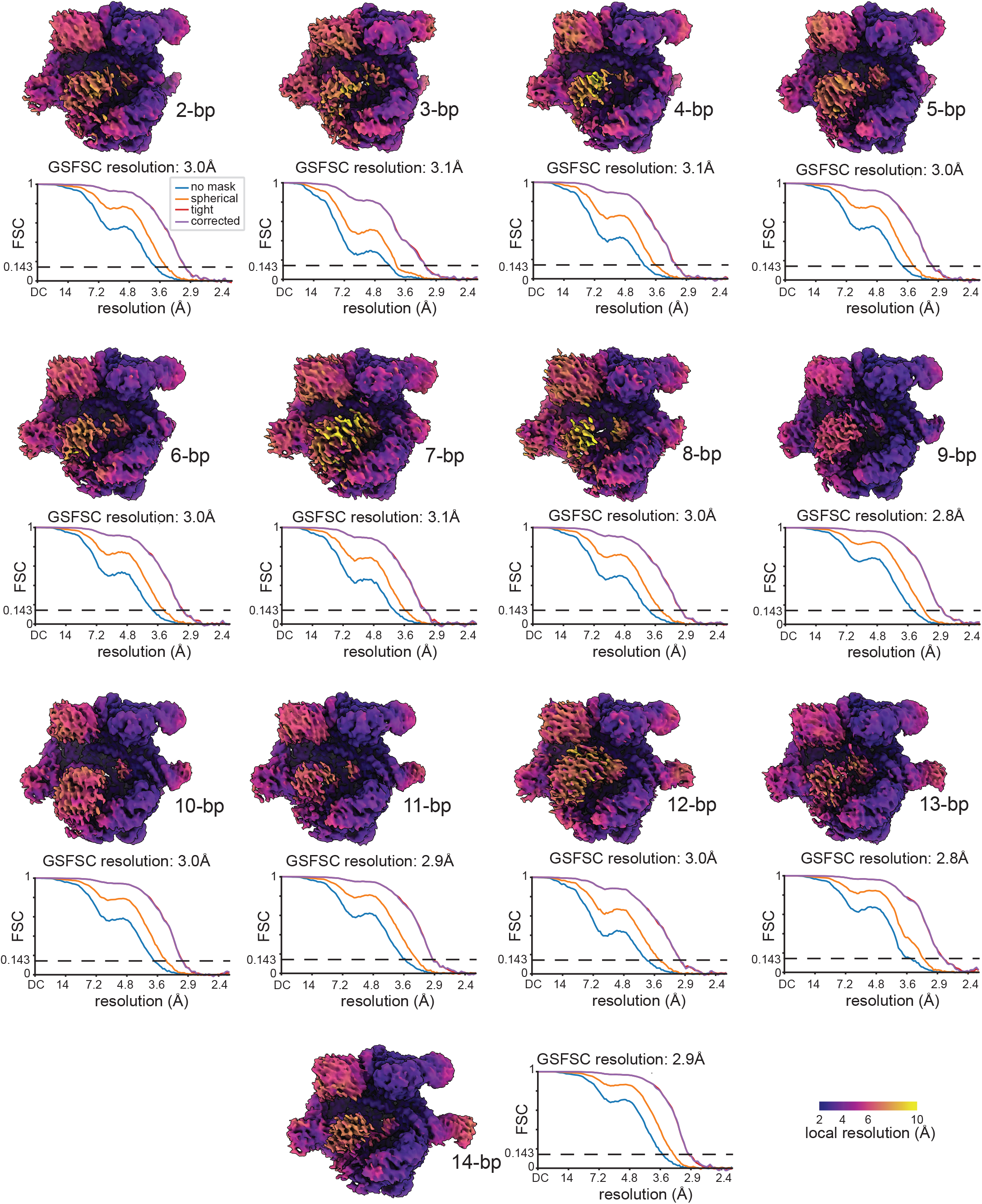
Individual datasets refine to high resolution despite non-encoded heterogeneity. Local resolution maps, colored according to the key (lower right) and Fourier shell correlation (FSC) plots from refinement of individual datasets (see Methods), as computed with various masks in cryoSPARC, are depicted for each of the 13 individual datasets. Resolution reported at 0.143 (dashed line) FSC criteria using a cryoSPARC “corrected” mask.

**Supplementary Figure 4.**
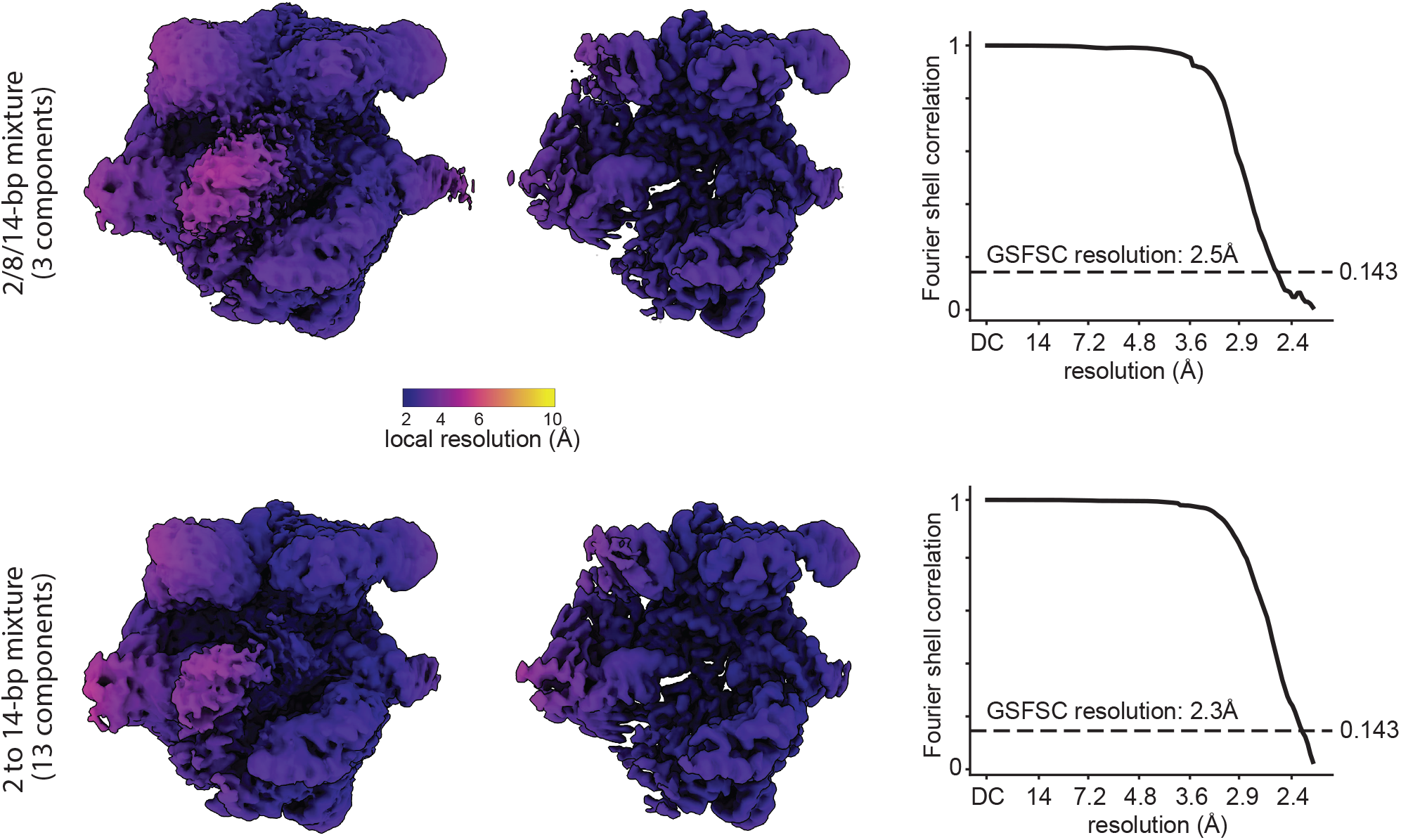
Consensus refinements on mixed particle stacks display substantial non-encoded heterogeneity. Local resolution maps from reconstructions of the 2/8/14-bp mixed particle stack (top) and the 2-14 bp mixed particle stack (bottom), colored according to the key. Surfaces from unsharpened maps shown at permissive (left) or restrictive (middle) contour levels. Fourier shell correlation (FSC) plots (right), computed in RELION, are shown with GS-FSC resolution reported at the 0.143 criteria (dashed line).

**Supplementary Figure 5.**
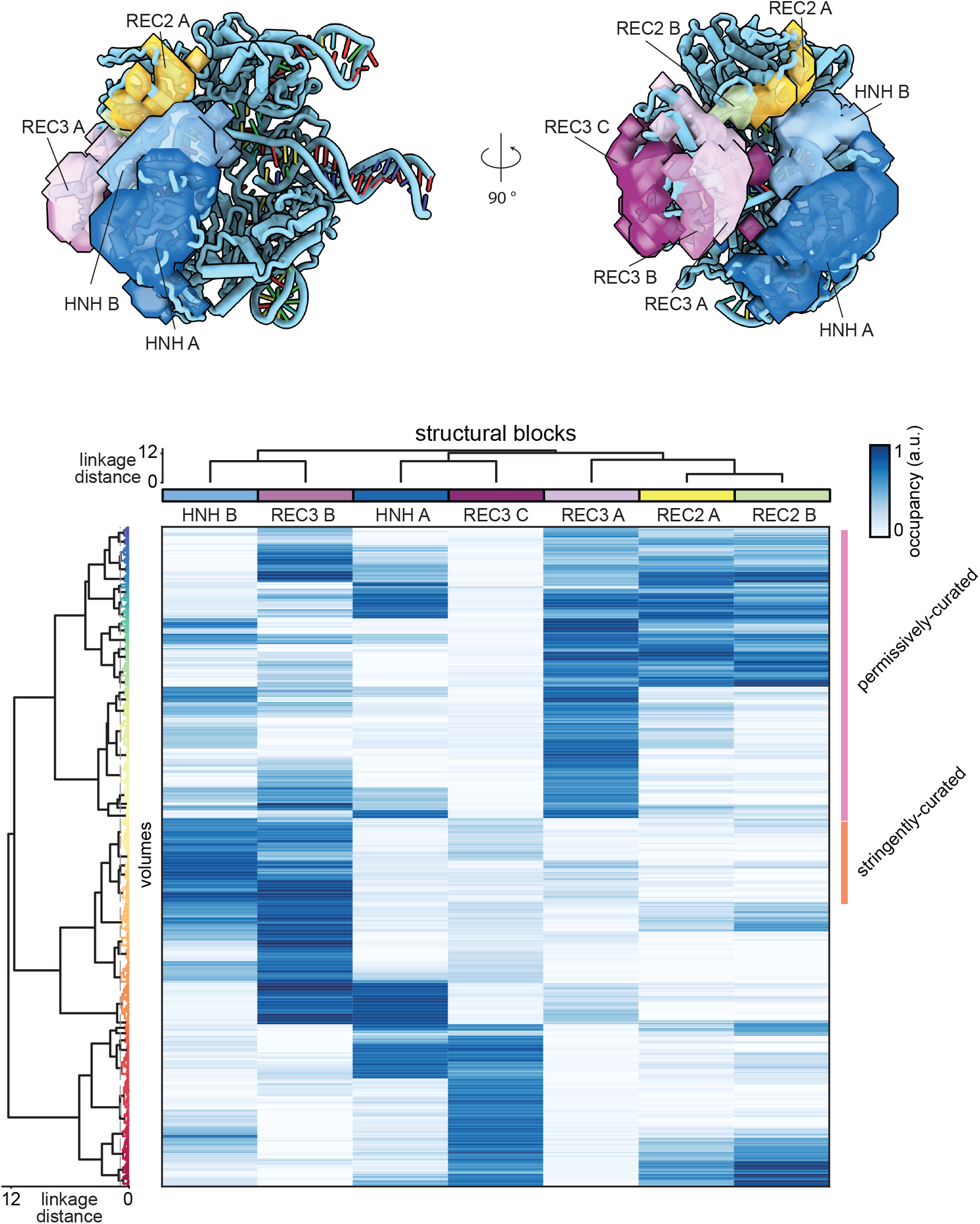
MAVEn-guided hierarchical clustering partitions a structural ensemble derived from deep classification into 48 classes. **(A)** SIREn structural blocks (semi-transparent surfaces) overlaid on atomic model of the dCas9•sgRNA•tDNA holocomplex (ribbon representation). Blocks are labeled according to dCas9 domain architecture (*i*.*e*., REC2, REC3, HNH), with alternative conformations distinguished as A-C. **(B)** Heatmap of normalized occupancies for eight selected SIREn blocks (labeled columns) across 500 deep classification volumes (rows). Structural blocks are colored as in (A). The dashed gray vertical line embedded in the dendrogram on the left indicates the distance threshold used to define classes. Colored bars on the right indicate classes pooled to generate curated subsets: pink for the permissively curated subset and orange for the stringently curated subset.

**Supplementary Figure 6.**
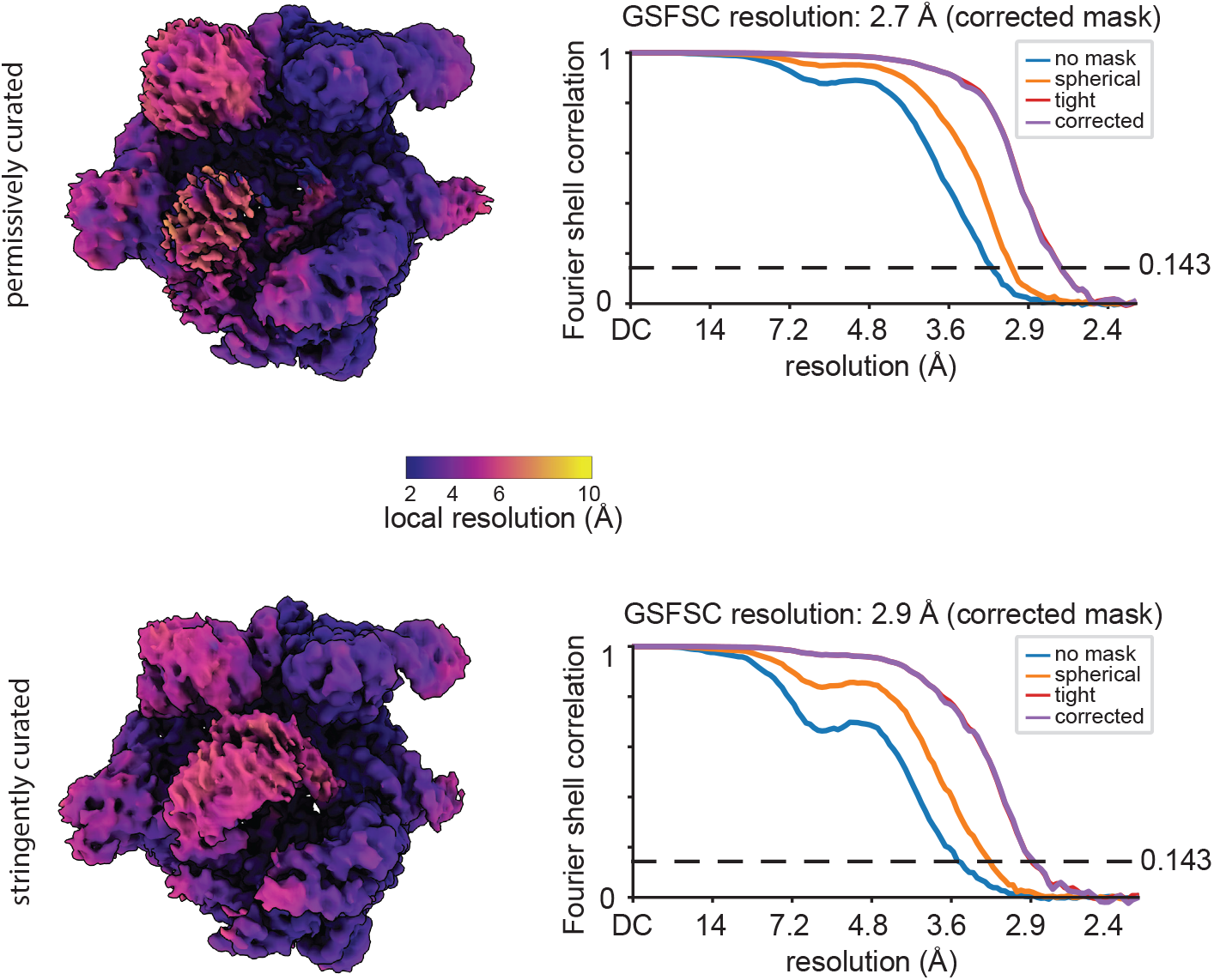
Curated subsets capture variable features at high resolution. Local resolution maps, colored following key, and FSC plots for permissively (top) and stringently (bottom) curated particle subsets.

**Supplementary Figure 7.**
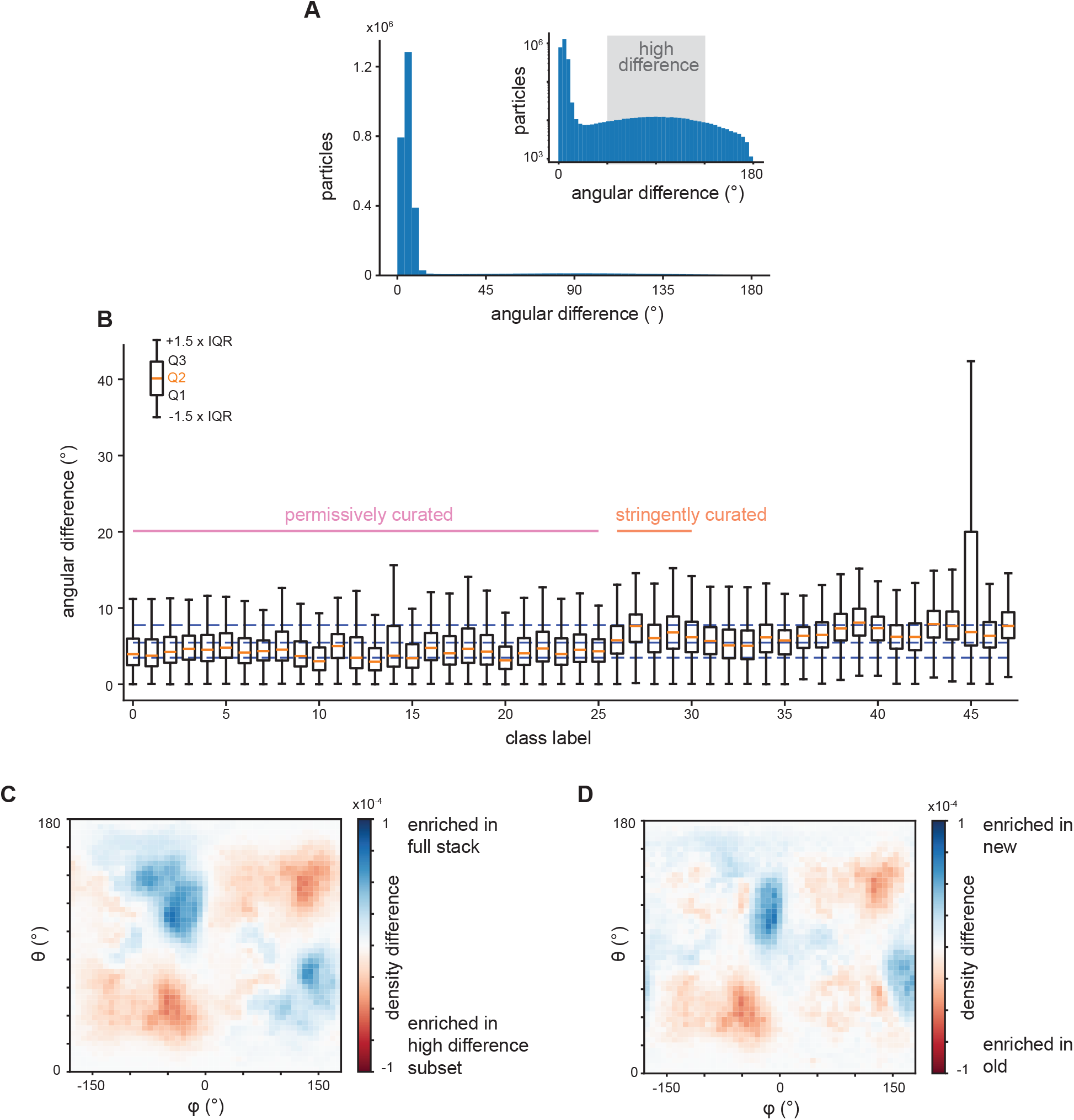
Deep classification refines pose estimates for a small subset of particles. **(A)** Distribution of angular differences between poses estimated by consensus refinement and the deep classification and pooled refinement approach for each particle (see Methods). Inset shows the same distribution with a log-scaled y-axis. Particles categorized as ‘high difference’ are indicated in grey. **(B)** Angular difference distributions conditioned on class label from the deep classification and pooled refinement approach. Each class label corresponds to a single pooled refinement. Classes that were combined to generate curated subsets are indicated. Dashed blue lines indicated 25^th^, 50^th^, and 75^th^ (Q1, Q2, Q3) percentiles across all particles. **(C)** Difference in consensus refinement pose distributions for the full particle stack and the high-difference subset. **(D)** Difference in pose distributions for the high-difference subset of particles, comparing poses from deep classification and pooled refinement (new) to poses from consensus refinement (old).

